# Genome-wide identification of genetic requirements of *Pseudomonas aeruginosa* PAO1 for rat cardiomyocyte (H9C2) infection by insertion sequencing

**DOI:** 10.1101/2021.03.03.433694

**Authors:** Jothi Ranjani, Ramamoorthy Sivakumar, Paramasamy Gunasekaran, Jeyaprakash Rajendhran

## Abstract

*Pseudomonas aeruginosa* is the major infectious agent among Gram-negative bacteria which causes both acute and chronic infections without any tissue specificity. Infections due to *P. aeruginosa* are hard to treat, as it entails various strategies like virulence factors synthesis, drug efflux systems & resistance and protein secretion systems during pathogenesis. Despite extensive research in *Pseudomonas* pathogenesis, novel drug targets and potential therapeutic strategies are inevitable. In this study, we investigated the genetic requirements of *P.aeruginosa* PAO1 for rat cardiomyocyte (H9C2) infection by insertion sequencing (INSeq). A mutant library comprising ~70,000 mutants of PAO1 was generated and the differentiated form of H9C2 cells (d-H9C2) was infected with the library. The infected d-H9C2 cells were maintained with antibiotic-protection and without any antibiotics in the growth media for 24 h. Subsequently, DNA library for INSeq was prepared, sequenced and fitness analysis was performed. A-One hundred and thirteen mutants were negatively selected in the infection condition with antibiotic-protection, whereas 143 mutants were negatively selected in antibiotic-free condition. Surprisingly, a higher number of mutants showed enriched fitness than the mutants of reduced fitness during the infection. We demonstrated that the genes associated with flagella and T3SS are important for adhesion and invasion of cardiomyocytes, while pili and proteases are conditionally essential during host cell lysis.

**Take away**

✓ **Fitness of *P.aeruginosa* mutants were analyzed during cardiomyocyte infection**
✓ **Genes involve amino acid transport & metabolism and signal transduction are important during intracellular lifestyle**
✓ **OMVs play a crucial role during infection and pathogenesis**
✓ **Flagella and T3SS are conditionally essential for adhesion and invasion, whereas pili and proteases are conditionally essential during host cell lysis**

## 1. Introduction

*Pseudomonas aeruginosa* is a Gram-negative, ubiquitous, and opportunistic pathogen responsible for chronic and acute infections. *P. aeruginosa* plays a critical role in chronic conditions such as cystic fibrosis, pneumonia, wound infections, chronic otitis media, chronic bacterial prostatitis, medical device-related infections and atherosclerotic carotid arterial plaques (Tolker-Nielsen, 2014 and Lanter *et al.*, 2014). Immunocompromised patients are highly susceptible to *P. aeruginosa* infections with consequent extensive damage to the organs; the most threatening pathogen in cystic fibrosis patients (Balasubramanian & Mathee, 2009) and the common source of blood-stream infections in cancer patients (Bos *et al.*, 2013). A plethora of researchers contributed their efforts towards understanding the pathogenesis mechanisms during *Pseudomonas* infections. Diverse approaches such as transcriptomics, proteomics, metabolomics, secretomics, high-throughput sequencing are employed in elucidating the pathogenesis mechanisms especially the host-pathogen interactions. Many efforts have been made to understand the host responses and pathogenesis especially in lung-related infections associated with cystic fibrosis. Here we demonstrated the conditional essentiality of the *Pseudomonas* genes during cardiomyocyte infection by employing insertion sequencing (INSeq).

INSeq is a reliable technique, which is a combination of mutagenesis and high throughput sequencing to screen a larger number of mutants with the relatively shortest duration. It is a robust and powerful technique for the rapid correlation of genotype to phenotype in a wide range of bacterial species. Based on transposon mutagenesis, various high throughput sequencing methods such as TraDIS, HITS, Tn-Seq, Tn-Seq Circle and INSeq have been developed. A well-established protocol for transposon library-based sequencing can be found elsewhere (Goodman *et al.*, 2011 and van opijnen *et al.*, 2010). Mutagenesis based sequencing has been performed in *Pseudomonas aeruginosa, Acinetobacter baumannii, Porphyromonas gingivalis, E coli*, *Streptococcus pneumonia, Moraxella catarrhalis*, *Bacteroides thetaiotaomicron*, *Streptococcus pyogenes*, etc., (Lee *et al.*, 2015, Gallagher *et al.*, 2015, Klein *et al.*, 2015, Shan *et al.*, 2015, Verhagen *et al.*, 2014, de Vries *et al.*, 2014, Goodman *et al.*, 2009, and Le Breton *et al.*, 2015). Earlier studies in host-pathogen interactions employed transposon library-based sequencing to identify the conditionally essential genes of the pathogen in the host environment. For instance, Turner *et al.*, (2014) demonstrated the genetic requirements of *P. aeruginosa* for acute burn wound and chronic infection by combining the RNA sequencing and insertion sequencing in a murine model. They found that the mutant fitness is not correlated with the expression of a gene *in vivo* except metabolic genes. Chemotactic motility and virulence factors are essential for acute burn wound infection but not for chronic wound infections. They concluded that the fitness of *P. aeruginosa* during infections is determined by the host physiology like the immune system (Turner *et al.*, 2014). The same group investigated the essential genes for survival in cystic fibrosis sputum (Turner *et al.*, 2015). Skurnik *et al.*, (2013) attempted to identify the conditionally essential genes of *P. aeruginosa* PA14 for mucosal colonization and systemic dissemination by INSeq. Gallagher *et al.*, (2011) identified the genes associated with intrinsic resistance to tobramycin in *P. aeruginosa*. Poulsen *et al.*, (2019) defined a core genome of *P. aeruginosa* and proposed novel drug targets. They identified 321 core essential genes from clinical isolates grown in analogous growth conditions such as serum, sputum and urine. *P.aeruginosa* fitness during biofilm growth was demonstrated by Schinner *et al.*, (2020).

In this study, we attempted to unveil the genetic requirement of *P.aeuroginosa* PAO1 to infect the rat cardiomyocytes through high throughput insertion sequencing. We identified 113 mutants that were negatively selected and 301 mutants were positively selected in the infection condition with amikacin when compared to the control library. Conversely, in the infection condition without any antibiotics, 143 mutants were negatively selected and 128 genes were positively selected. Our data substantiate the fact that the flagella and T3SS are conditionally essential for adhesion and invasion whereas the pili and proteases are essential for host cell lysis.

## 2. Materials and Methods

### 2.1. Bacterial strains and plasmids

*Pseudomonas aeruginosa* PAO1 was routinely cultured in Luria-Bertani (LB) broth at 37 °C. Bacteria at exponential phase were used for infecting the host cells. *E. coli* Sm10λpir harboring pSAM_BT20 plasmid was maintained in LB broth with 100 μg/ml ampicillin. PAO1 mutant strains were procured from *P. aeruginosa* mutant library, University of Washington and maintained in LB with 5 μg/ml tetracycline (Jacobs *et al.*, 2003 and Held *et al.*, 2012).

### 2.2. Cell line maintenance and differentiation

H9C2 cell line was obtained from the National Centre for Cell Science, Pune, India. Cells were cultured in Dulbecco’s Modified Eagle’s Medium (DMEM) F12 Ham supplemented with 10% Fetal Bovine Serum (FBS), 100 U/ml of penicillin, 100 μg/ml of streptomycin and 2.5 μg/ml of amphotericin B in a humidified incubator with 5% CO_2_ at 37°C. Cardiomyoblast differentiation was induced as previously described (Ranjani *et al.*, 2015). Briefly, H9C2 cells were seeded in 75 cm^2^ tissue culture flasks at a density of 3.5×10^4^ cells in the growth medium. At 50-60% of confluence, high serum media was replaced by low serum media (1% FBS) and incubated for 4 days in the dark with a daily supplement of 1 μM all-trans retinoic acid. On the fifth day of treatment, differentiated cells were trypsinized with 0.025% trypsin-EDTA solution and seeded for further experiments.

### 2.3. Generation of PAO1 mutant library

*P. aeruginosa* PAO1 mutant library was generated according to Skurnik *et al.*, (2013) with few modifications. Briefly, an overnight culture of *E. coli* Sm10λpir pSAM_BT20 (Sivakumar *et al.*, 2019) and *P. aeruginosa* PAO1 were mixed and centrifuged. The pellet was washed in LB, resuspended in 100 μl of LB, spotted on pre-warmed LB agar plates and incubated at 37 °C for 3 h. The conjugation spots were suspended in LB and plated on LB agar containing 25 μg/ml of irgasan and 40 μg/ml of gentamicin. After 16 h of incubation, the colonies were collected with a cell scraper, suspended in PBS with 15 % glycerol and stored at −80 °C until further use. Approximately 70,000 mutants were generated and the transposition was confirmed by amplifying the gentamicin cassette and transposase using the primers GENF-GENR and TRANSF-TRANSR respectively (Datafile-1).

### 2.4. d-H9C2 infection with mutant library

Infection assays were performed according to our previous study (Ranjani *et al.*, 2015). d-H9C2 cells were infected with the mutant library at 1:25 MOI for 1 h. After the infection period, the d-H9C2 cells were incubated with amikacin containing growth media for 1h. Further, the cells were washed and maintained in two different conditions such as in-presence and absence of amikacin in the growth media. At 24 h of post-infection, infected cells were harvested, and genomic DNA was isolated (Qiagen DNeasy blood and tissue kit).

### 2.5. Library preparation for insertion sequencing

DNA library was prepared as previously mentioned (Sivakumar *et al.*, 2019), Briefly, insertion junctions were amplified using BiosamA primer followed by clean up and streptavidin bead binding. The second strand was synthesized using random hexamers (Roche, USA) followed by MmeI digestion. After digestion, adapters were ligated, amplified and size selected. Sequencing was performed in Ion Torrent personal genome machine using a 318v2 chip and a 200 bp sequencing chemistry.

### 2.6. Data analysis

Adapters were trimmed from the raw reads using Cutadapt, and the reads were split based on barcodes using FastX barcode splitter. The analysis was performed using the online software ESSENTIALS (http://bamics2.cmbi.ru.nl/websoftware/essentials) (Zomer *et al.*, 2012) by choosing *P. aeruginosa* PAO1 (NC_002516) as the reference genome.

### 2.7. Live/dead cell assay

The cytotoxic effect of the wild-type and mutant strains were analyzed by live/dead cell assay using the High Content Screening (HCS) System (Operetta, Perkin Elmer, USA). At 12 h of post-infection, media was aspirated carefully, and cells were stained with Hoechst 33342 and PI (Invitrogen, USA). Images were captured, and analysis was performed in Harmony software. Twelve fields were analyzed per condition in triplicate, and the results were given as mean values.

### 2.8. CFU enumeration

d-H9C2 cells were infected with wild type and mutants, harvested by scrapping at 12 h of post-infection periods and lysed with 0.1% Triton X-100 in PBS. The lysate was plated onto antibiotic-free King’s B media after serial dilution.

### 2.9. Image process and statistical analysis

HCS Images were processed using ImageJ 1.49 (Schneider *et al.*, 2012). All the experiments were performed in triplicates, and the data represented in bar charts are the means of three independent experiments with standard deviation. Statistical analyses were performed using GraphPad QuickCalcs.

## 3. Results and discussion

### 3.1. Unique insertions and essential genes

Approximately, 7.5 million reads were obtained from the triplicate samples of input library (DMEM), and 24 h infected cells in two different conditions. Adapter sequences were trimmed, reads were binned based on the barcodes, and analyzed using ESSENTIALS software. About 65536 unique insertions were found in both strands of the genome. Based on the density plot, 471 genes were identified as essential for PAO1 to survive in the DMEM medium. Among 471, two hundred and sixty-eight genes were already reported as the core essential genome of *P.aeruginosa* by Poulsen *et al.*, (2019). The orthologs of PAO1 for the essential genes reported in PA14 (Skurnik *et al.*, 2013 and Poulsen *et al.*, 2019) were found from the *Pseudomonas* database (Winsor *et al.*, 2016) and included in the comparison along with other studies (Liberati *et al.*, 2006, Turner *et al.*,2014 and Lee *et al.*, 2015). The intersection with all previous studies showed that about 293 genes identified as essential in the current study were reported in any one of the studies. We also found 89 essential genes as common-the core-set in all these studies (Figure 1). Genes associated with ribosomal structure, cell wall/membrane/envelope and their biogenesis were found to be the major categories of the essential genes followed by various metabolite transport and metabolism. The ribosome plays a fundamental role in the bacteria and hence many existing antibiotics bind to ribosome for inhibiting several downstream processes. Moreover, the essential genes of the pathogen are the potential drug targets and hence, further exploration of these essential genes particularly the core-set of essential genes will lead to promising candidate drug targets. A list of identified essential genes and COG categories is provided in Datafile-2.

**Figure 1.**
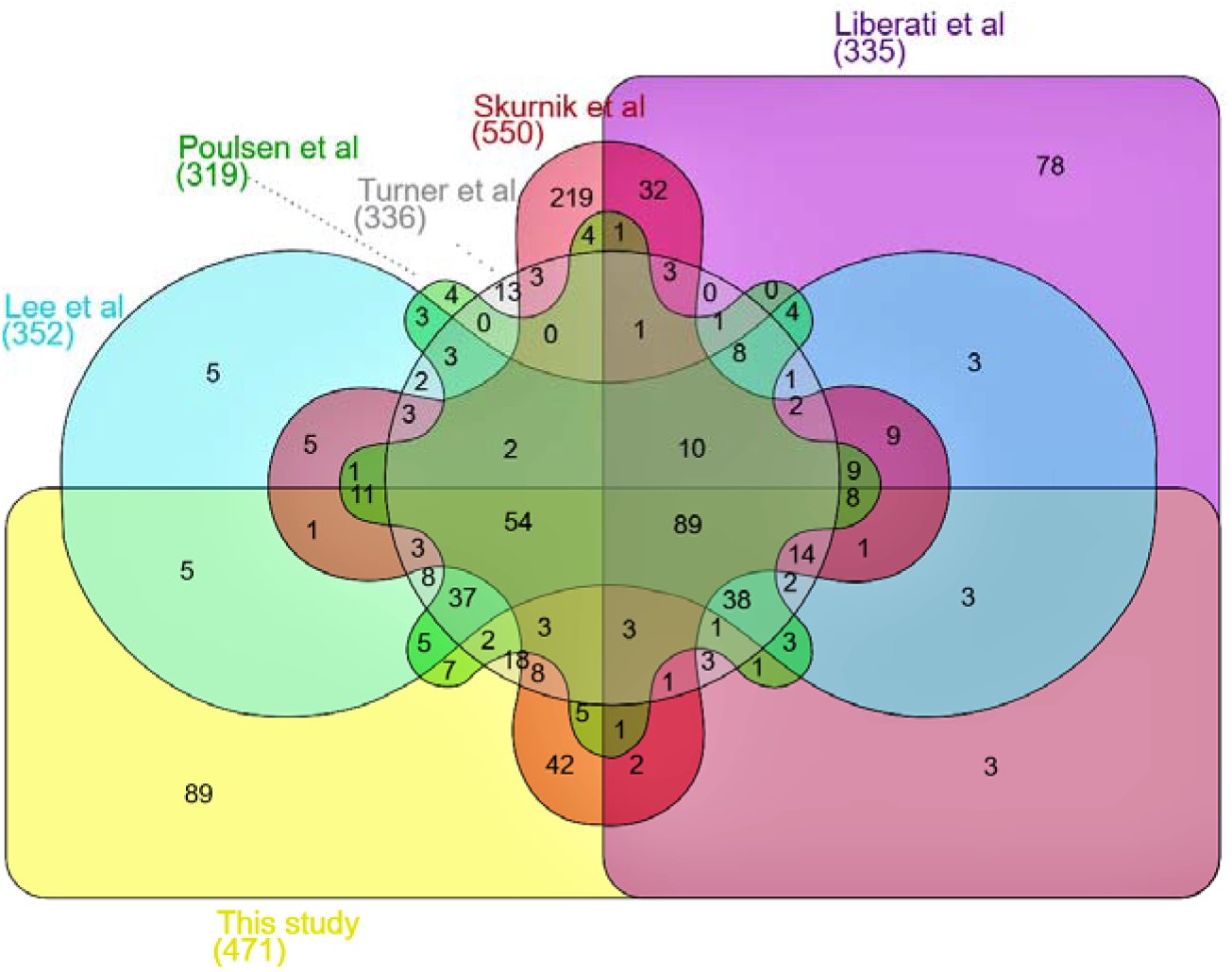
The intersection of the identified essential genes with previous studies: Among 471 essential genes, 89 genes are identified as essential in all the previous studies (Core-set) and 293 genes were reported already in any one of these studies. Venn diagram was generated using Interactivenn (Heberle *et al.*, 2015).

### 3.2. Genetic requirements of PAO1 for cardiomyocyte infection

Although many studies revealed essential genes of *Pseudomonas* for infection, none of them were at the cellular level, particularly at the intracellular condition. Here we maintained the infected cardiomyocytes with antibiotic-protection and without antibiotics. During antibiotic-protection, pathogens that are actively released from the host as well as during host lysis will be killed by the extracellular antibiotic–amikacin. Since mammalian cells are impermeable to amikacin, internalized pathogens will be protected. On other hand, withdrawal of antibiotics after rigorously washing the infected cells aids the maintenance and growth of the whole population of the pathogen. Hence, post-infection in the presence of antibiotics helps specifically to study the intracellular population and without antibiotics aid to analyze the entire population including released from the host cells.

#### 3.2.1. Conditionally essential genes of PAO1 for cardiomyocyte adhesion and infection

We obtained 113 negatively selected mutants and 301 positively selected mutants in antibiotic-protection condition after infection (Figure 2a and 2b). Surprisingly, a higher number of mutants showed enriched fitness in this condition than the negatively selected mutants (Datafile-3). Among negatively selected mutants, PA1561, PA5205, PA0928, PA5535, PA2197, PA0271, PA2559 and PA5045 showed more than four-fold changes particularly mutant of PA1561 had −63.381-fold difference. PA1561 encodes aerotaxis receptor Aer, PA5205 is a conserved hypothetical MarC-related protein, PA0928 encodes GacS, PA5535, PA2197, PA2559 and PA0271 are hypothetical proteins, and PA5045 is penicillin-binding protein 1A PonA. The fitness of negatively selected mutants is illustrated in Table 1.

**Figure 2.**
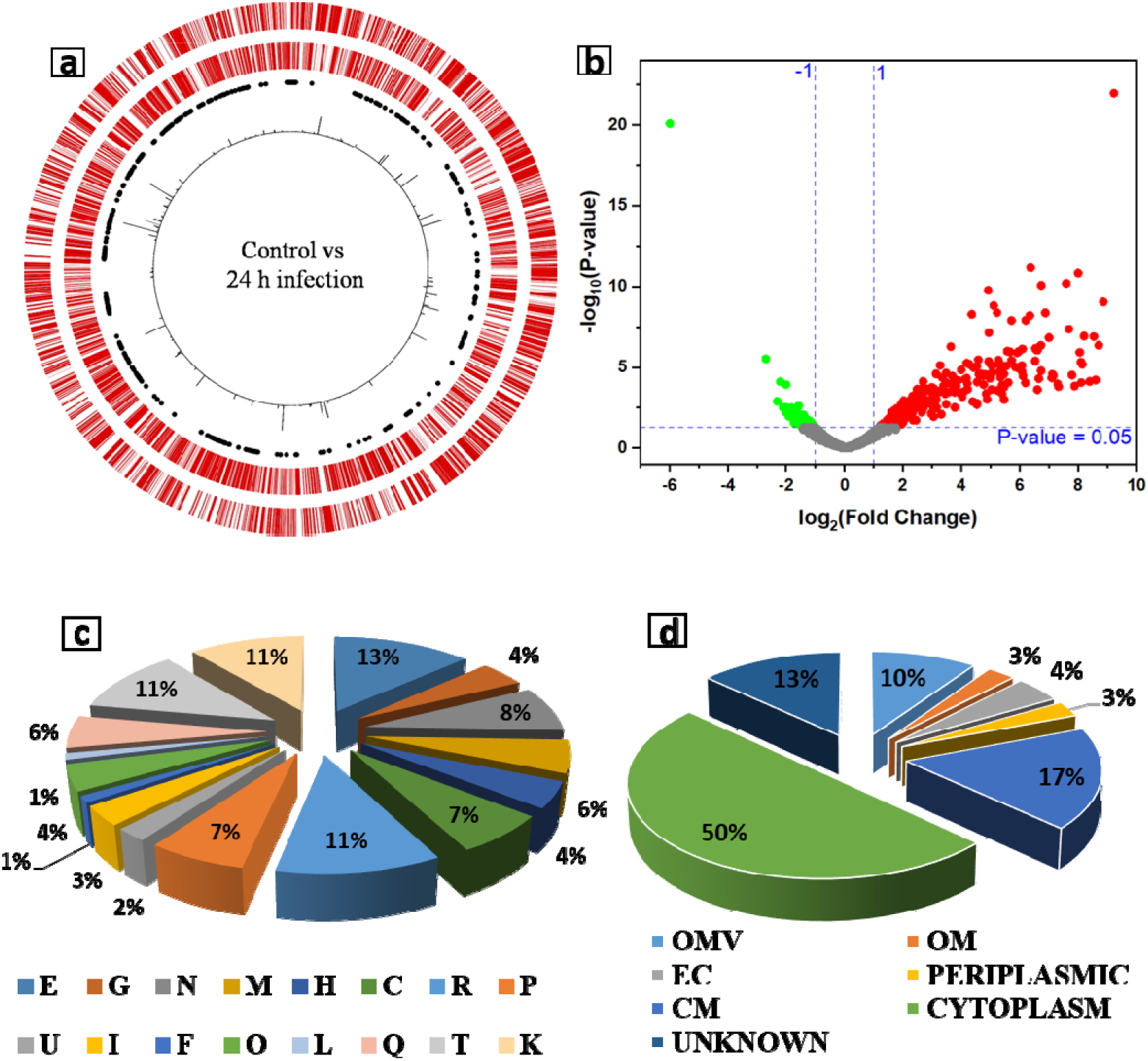
**a**. Representation of the fold change occurrence in the mutants during infection condition with amikacin. From the outermost ring to innermost ring: genes on the positive strand, genes on the negative strand, 471 identified essential genes, fold change occurrence in the mutants (outward line-positive fold change, inward line-negative fold change). The circular map was drawn using CiVi, a web-based tool (Overmars *et al.*, 2015). **b**. Volcano plot representing the fitness of the mutants. **c.** COG categories of conditionally essential genes for d-H9C2 infection. **d**. Categories of conditionally essential genes based on localization. Percentage of distribution in a various subcellular location: OMV: Outer membrane vesicles, OM: Outer membrane, EC: Extracellular, CM: Cytoplasmic membrane

**Table 1.**
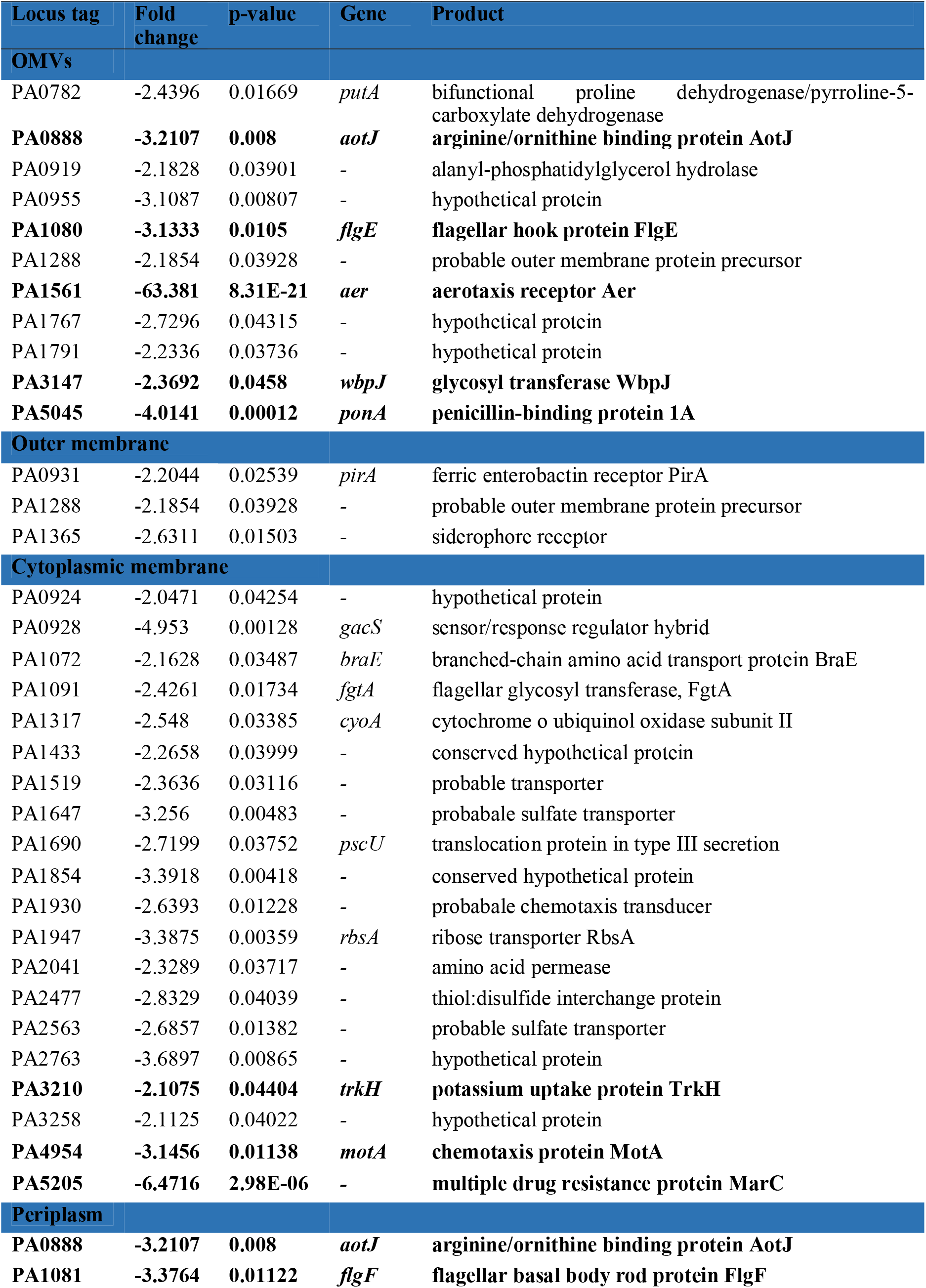
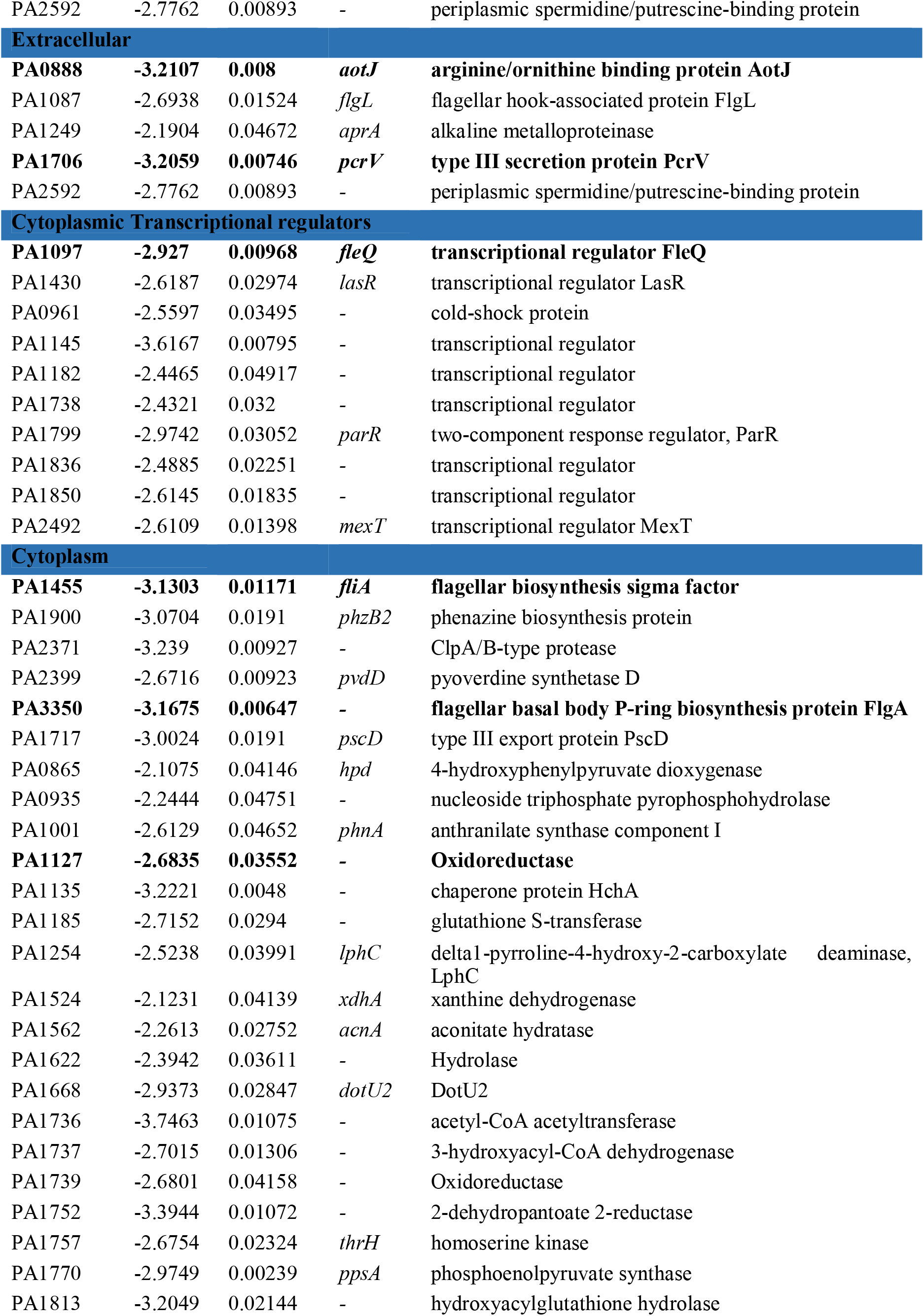
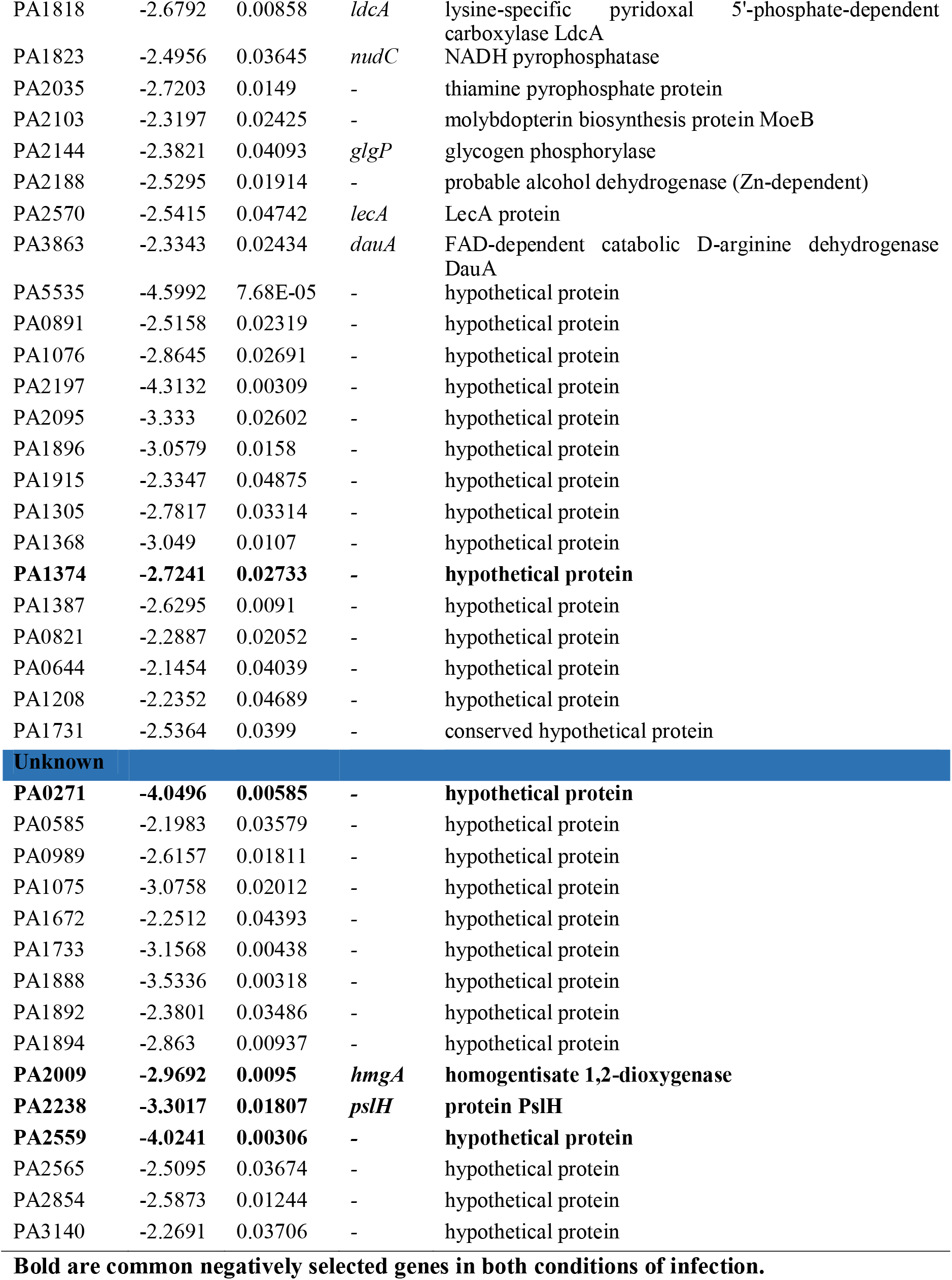
Conditionally essential genes of PAO1 in antibiotic-protection condition after d-H9C2 infection.

Conditionally essential (negatively-selected) genes during this condition belong to various COG categories (Figure 2c). Amino acid transport and metabolism (13%) is the major class followed by Signal transduction mechanisms and Transcription (11%). About 8% of genes belong to cell motility, and 7% of genes belong to energy production & conservation, and Inorganic ion transport & metabolism. Fifty percent of the population localizes in the cytoplasm. About 17% localize in the cytoplasmic membrane, 10% are outer membrane vesicle (OMV), 4% are extracellular, and 3% are outer membrane and periplasm (Figure 2d). A fifteen percent of the conditionally essential genes are already known virulence factors. Of these virulence factors, many genes are associated with flagella such as *flgE, flgF, flgL, fliA, fgtA*, PA3350, and *motA*. *pcrV, pscD,* and *pscU* belong to type-three secretion system and a type VI secretion system gene PA2371 – *clpV3*. *gacS, lasR, wbpJ, pvdD* and *aprA* are other virulence factors found to be conditionally essential during infection. The fitness level of flagella and T3SS associated mutants suggests their crucial role in adhesion and infection. The flagellum is a complex organelle and an assembly of more than 20 proteins (Haiko *et al.*, 2013). More than 50 genes involved in flagellar biosynthesis and function. Flagella play a vital role in pathogenesis through their motility and chemotaxis properties. Besides its role in motility, it also acts as the ligand for many eukaryotic receptors such as asialo ganglio-N-tetraosylceramide (asialo-GM1), toll-like receptor 2 (TLR2), etc., (Feldman *et al.*, 1998). The whole flagellum aids a significant role in the adhesion of host cells and acts as an antigen for adaptive immunity (Haiko *et al.*, 2013). The importance of flagellar components in host-pathogen interactions was demonstrated in many studies (Verma *et al.*, 2006, Shen *et al.*, 2017, Jyot *et al.*, 2007, Zhang *et al.*, 2007 and Andersen-Nissen *et al.*, 2005). Reinforcing the previous studies, our INSeq data also evidenced the significance of flagella specifically during adhesion and infection of cardiomyocytes.

T3SS is the major virulence factor of *P. aeruginosa,* and it is active during acute infections (Hauser *et al.*, 2002). Toxins or effectors are directly injected into the host cell through T3SS to hijack the cellular pathways with consequent cell death. T3SS is a syringe-like structure that is an assembly of over 20 proteins in three complexes called the basal body, the needle, and the translocon. We found *pcrV, pscD* and *pscU* are conditionally essential for cardiomyocyte infection in the presence of amikacin. *pcrV* – low calcium response locus protein V is the needle tip of the T3S translocator and is required for the appropriate assembly of the injector (Sato *et al.*, 2011). *pcrV* also regulates the effector translocation and an important protective antigen against T3SS (Frank *et al.*, 2002 and Holder *et al.*, 2001). Hence, PcrV is a potential target of the therapeutic antibody (Thaden *et al.*, 2016). PscD and PscU are the components of the basal body where the former present in the inner membrane and the latter present in the cytoplasmic phase (Perdu *et al.*, 2015). PscU is an autoprotease and also a ruler protein that regulates the length of the needle (Bergeron *et al.*, 2016).

Around 37% of conditionally essential gene products belong to membrane proteins including OMVs, outer membrane, periplasmic and cytoplasmic membrane. Ten percent of the products localize in OMV. OMVs are the spherical vesicles bud off from the outer membrane filled with periplasmic contents. OMVs are the mediator of the bacterium to interact with various environments such as host niche, stress, etc., (Schwechheimer and Kuehn, 2015). Host cell-targeted delivery, protection from proteolytic dehydration, modulation of the immune response, bacterial effector secretion and distant delivery of proteins are the most significant benefits of OMVs. In a host niche, mostly OMVs are enriched with virulence factors that aid in the host cell invasion, destruction, immune system evasion and antibiotic resistance (Bonnington and Kuehn, 2014). PutA, AotJ, FlgE, Aer, WbpJ, and PonA are some gene-products localize in OMVs and the rest are hypothetical proteins. PutA is a bifunctional proline dehydrogenase/pyrroline-5-carboxylate dehydrogenase, AotJ- arginine/ornithine-binding protein, FlgE- flagellar hook protein, Aer- aerotaxis receptor, WbpJ- glycosyltransferase and PonA is penicillin-binding protein 1A. WbpJ and FlgE have known virulence factors, where the former involve in LPS O-antigen synthesis (Rocchetta *et al.*, 1999) and the latter is a flagellar component. PAO1 utilizes proline as the sole source of carbon and nitrogen by using the bifunctional PutA (Nakada *et al.*, 2002). AotJ belongs to the ABC transporter system which binds to arginine/ornithine. Everett *et al.*, (2017) demonstrated that the virulence of *Pseudomonas* is highly influenced by the presence of arginine in the host environment especially in vivo because arginine is a conditionally essential amino acid for the pathogen. Aer aids the bacterium to identify the specific oxygen and energy-rich niches (Amin *et al.*, 2007). *fleQ, lasR, mexT, parR*, PA0961, PA1145, PA1182, PA1738, PA1799, PA1836, and PA1850 are the conditionally essential transcriptional regulators during this cardiomyocyte infection condition. FleQ is the flagellar transcriptional regulator that highly influences flagellar biogenesis and its related operons (Dasgupta *et al.*, 2002). FleQ also regulates the expression of *pel* genes by acting as both the activator and repressor based on the availability of c-di-GMP (Baraquet *et al.*, 2012). LasR is the global regulator of many virulence factors and an important quorum-sensing receptor. Elastases, pyocyanin, and proteases are the virulence factors whose expression is highly regulated by LasR (Longo *et al.*, 2013). The role of other conditionally essential transcriptional regulators needs to be analyzed for understanding their potential function during infection.

#### 3.2.2. Conditionally essential genes of PAO1 for cardiomyocyte lysis/active release

We found 143 negatively selected mutants and 128 positively selected mutants, in the antibiotics-free condition after d-H9C2 infection (Figure 3a and 3b). About fifteen mutants showed more than four-fold reduced fitness, and nine genes showed more than four-fold increased fitness. Mutants of PA5471, PA0395, PAO271, PA4547, PA1561, PA1101, PA0766, PA1758, PA1374, PA1097, PA4398, PA3824, PA0375, PA4546, and PA0396 were shown reduced fitness. Table 2 shows the list of conditionally essential genes and their fold-change during d-H9C2 infection without amikacin and Datafile-3 is the list of positively selected mutants. Many genes involved in motility especially the adhesins such as *pilA, pilT, pilU, pilO, pilN, flgE, flgF, flhA, flhB, and motA* are found to be conditionally essential. Genes associated with signal transduction and transcription are the next major group of conditionally essential genes followed by nutrient transport systems specifically carbohydrate and amino acid (Figure 3c). A twenty-six percent of conditionally essential gene products for d-H9C2 infection without amikacin are localized in the cytoplasmic membrane. The majority of the gene products belong to the cytoplasm (39%), 10% are outer membrane vesicle, 3% are outer membrane, periplasm and extracellular. A sixteen percent of the population’s localization is unknown (Figure 3d).

**Figure 3.**
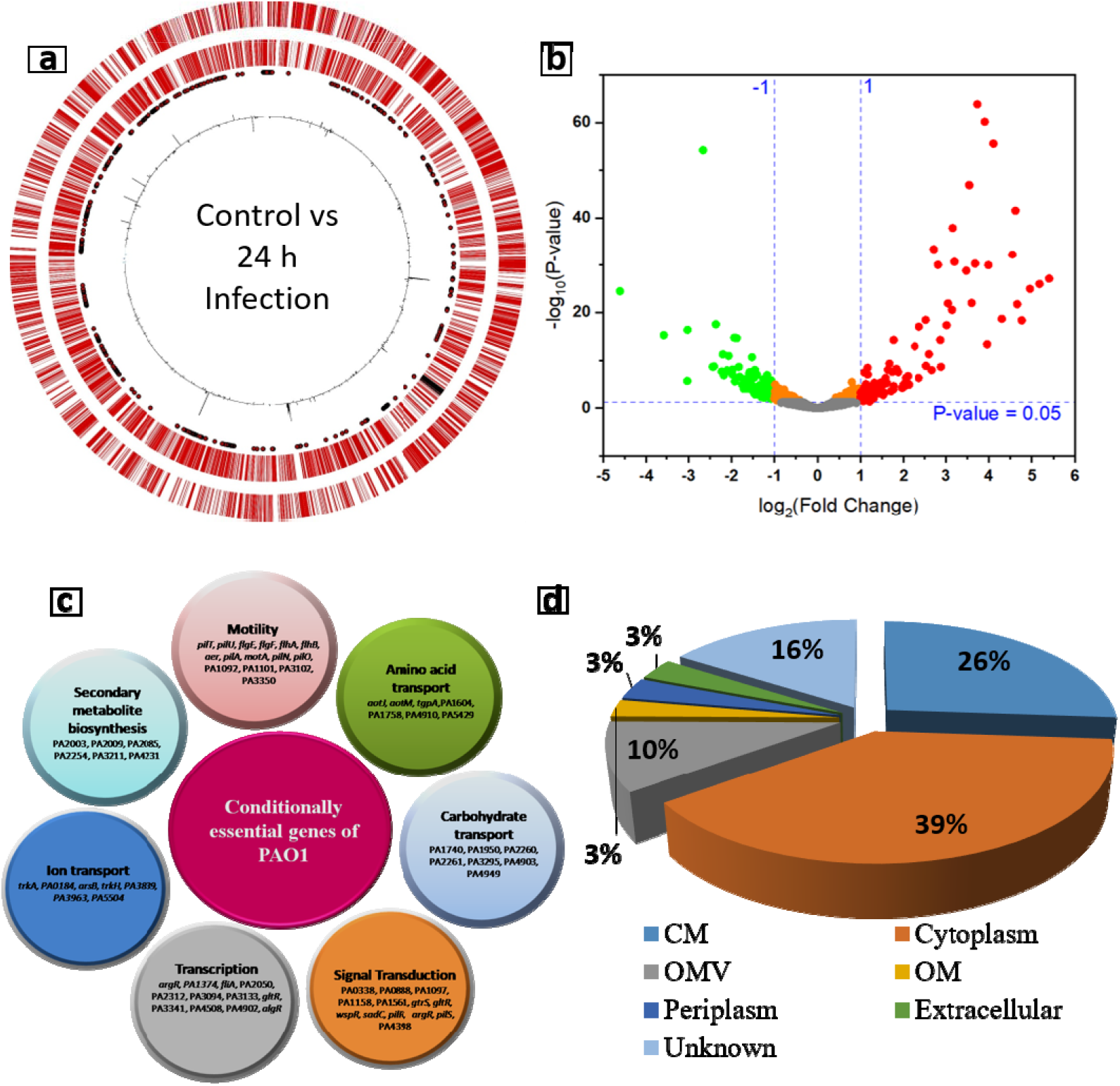
**a.** Representation of the fold change occurrence in the mutants in the infection condition without antibiotics. From the outermost ring to innermost ring: genes on the positive strand, genes on the negative strand, 471 identified essential genes, fold change occurrence in the mutants (outward line-positive fold change, inward line-negative fold change). **b**. Volcano plot representing the fitness of the mutants. **c.** Representative image of the major COG categories of conditionally essential genes during d-H9C2 infection without amikacin. **d.** Categories of conditionally essential gene products based on localization. Percentage of distribution in a various subcellular location: OMV: Outer membrane vesicles, OM: Outer membrane, EC: Extracellular, CM: Cytoplasmic membrane.

**Table 2.**
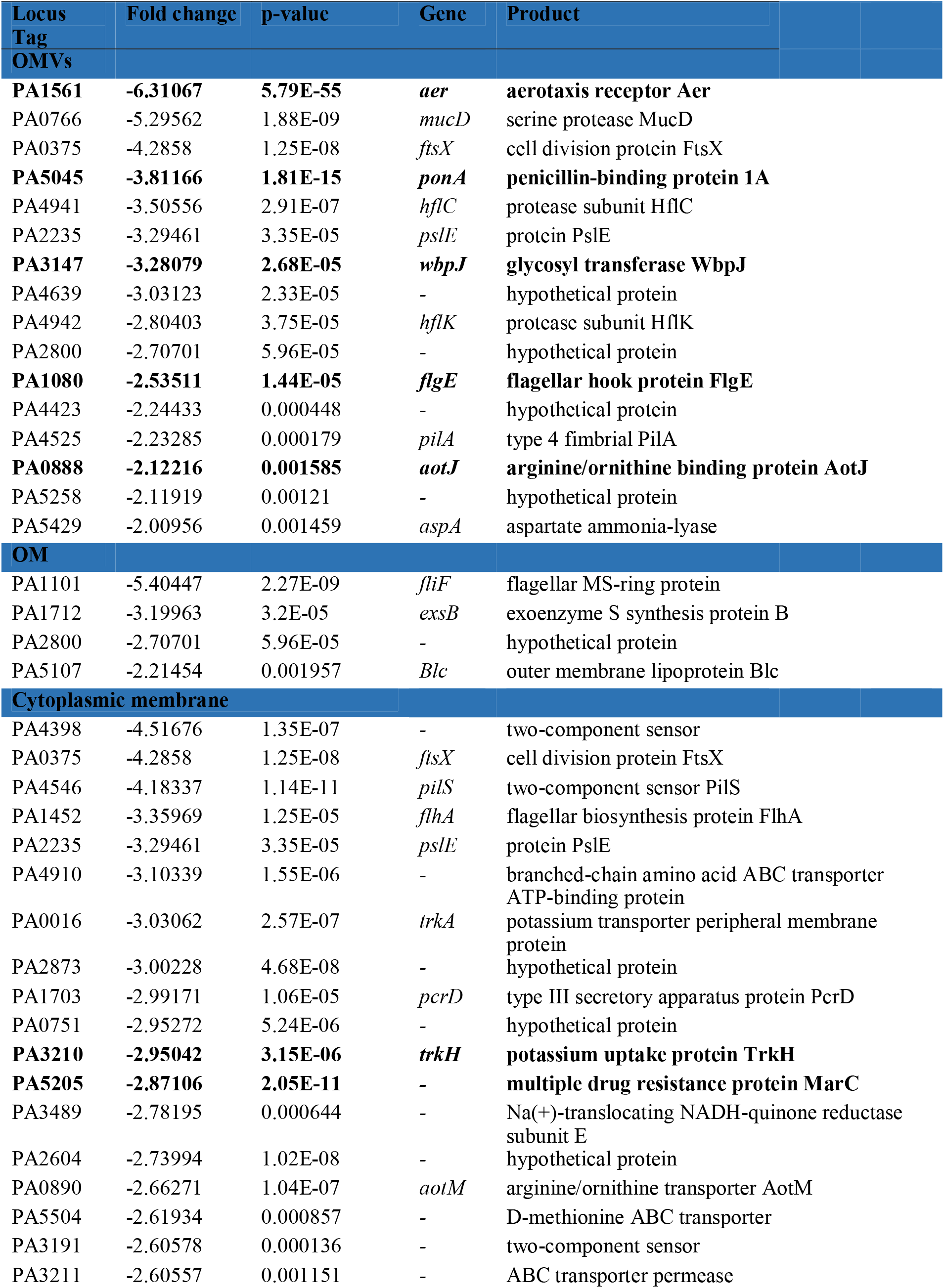
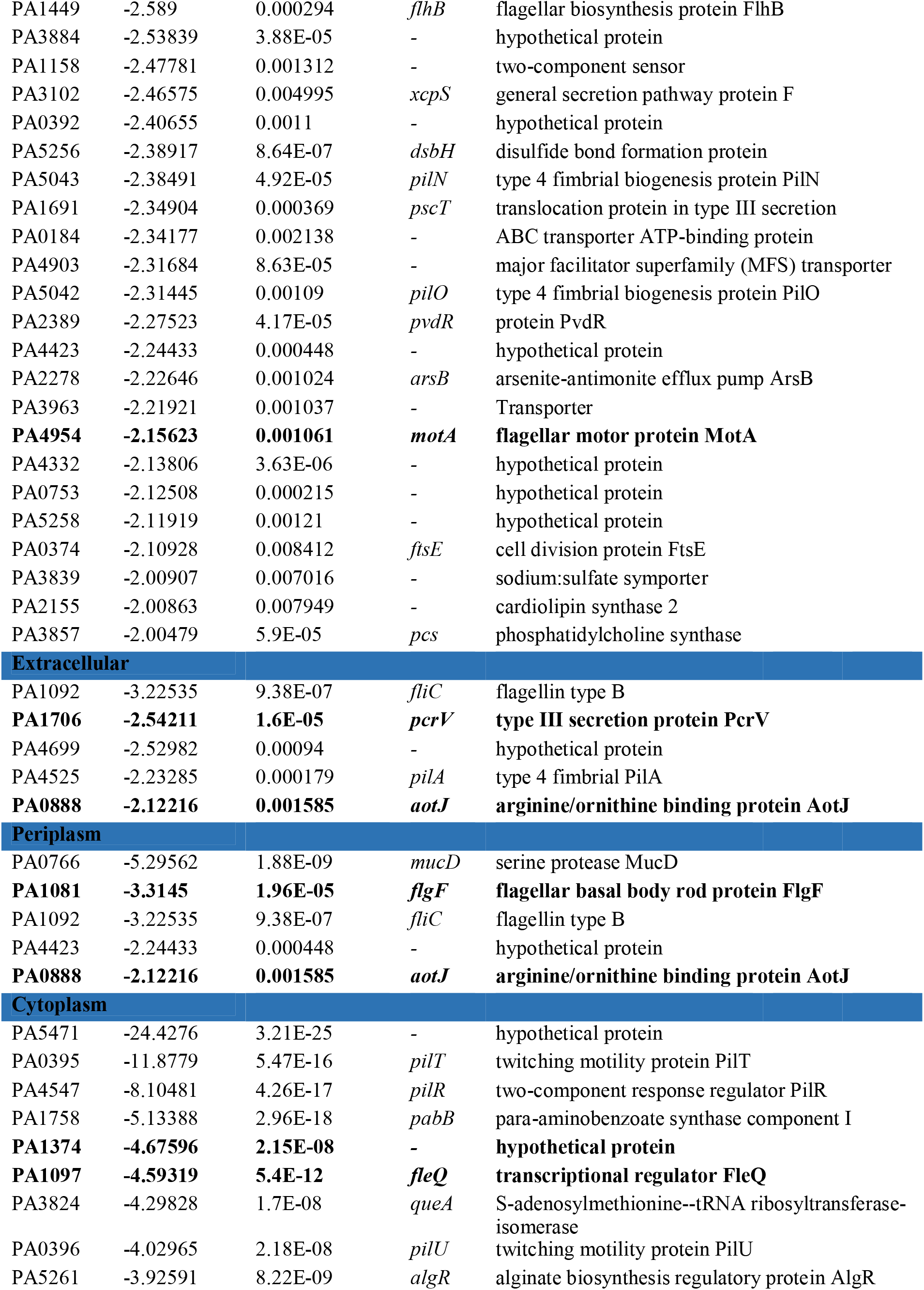
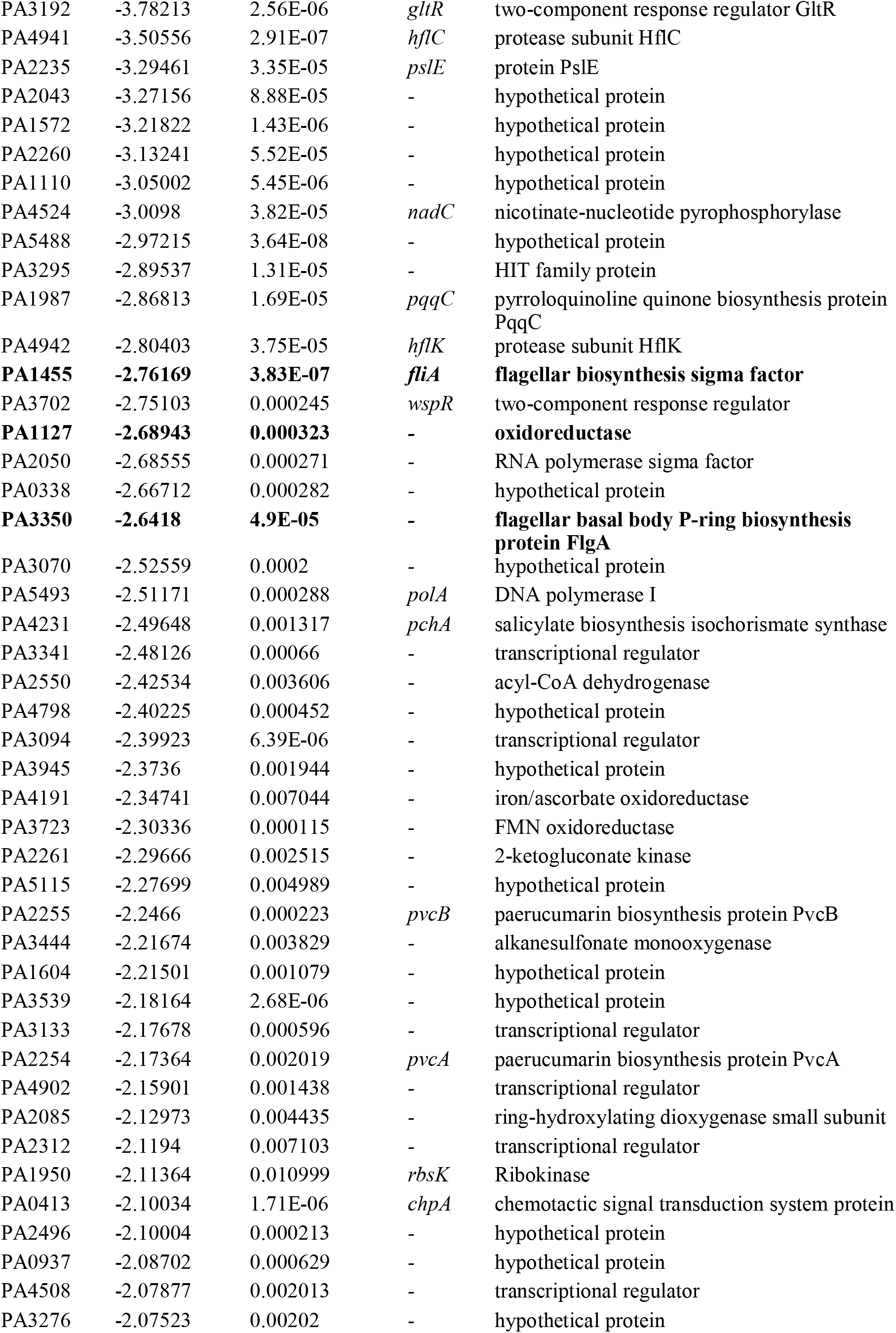
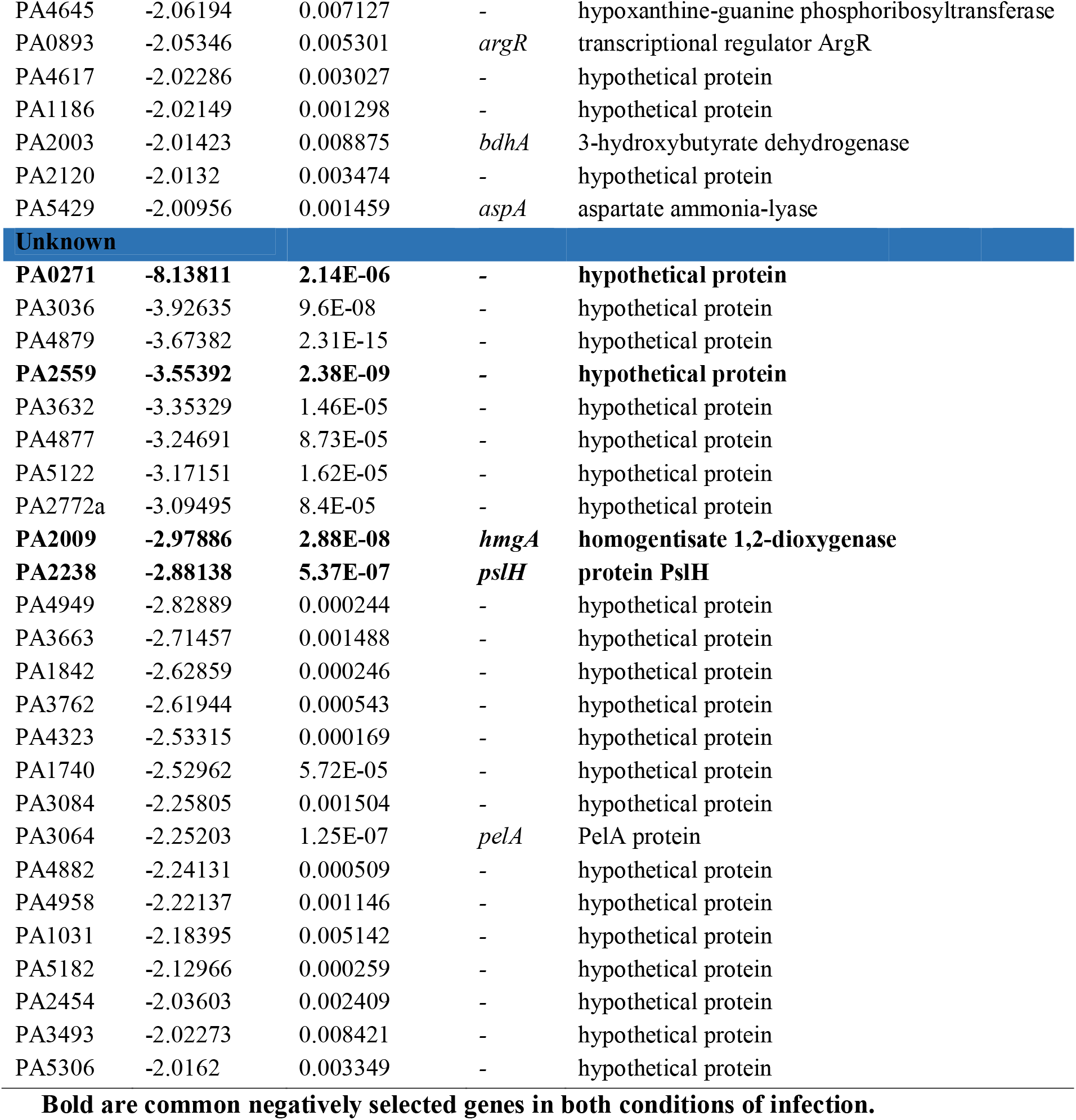
Conditionally essential genes of PAO1 in antibiotics-free condition after d-H9C2 infection.

| Locus Tag | Fold change | p-value | Gene | Product |
| --- | --- | --- | --- | --- |
| **OMVs** |  |  |  |  |
| **PA1561** | **-6.31067** | **5.79E-55** | *aer* | **aerotaxis receptor Aer** |
| PA0766 | -5.29562 | 1.88E-09 | *mucD* | serine protease MucD |
| PA0375 | -4.2858 | 1.25E-08 | *ftsX* | cell division protein FtsX |
| **PA5045** | **-3.81166** | **1.81E-15** | *ponA* | **penicillin-binding protein 1A** |
| PA4941 | -3.50556 | 2.91E-07 | *hflC* | protease subunit HflC |
| PA2235 | -3.29461 | 3.35E-05 | *pslE* | protein PslE |
| **PA3147** | **-3.28079** | **2.68E-05** | *wbpJ* | **glycosyl transferase WbpJ** |
| PA4639 | -3.03123 | 2.33E-05 | - | hypothetical protein |
| PA4942 | -2.80403 | 3.75E-05 | *hflK* | protease subunit HflK |
| PA2800 | -2.70701 | 5.96E-05 | - | hypothetical protein |
| **PA1080** | **-2.53511** | **1.44E-05** | *flgE* | **flagellar hook protein FlgE** |
| PA4423 | -2.24433 | 0.000448 | - | hypothetical protein |
| PA4525 | -2.23285 | 0.000179 | *pilA* | type 4 fimbrial PilA |
| **PA0888** | **-2.12216** | **0.001585** | *aotJ* | **arginine/ornithine binding protein AotJ** |
| PA5258 | -2.11919 | 0.00121 | - | hypothetical protein |
| PA5429 | -2.00956 | 0.001459 | *aspA* | aspartate ammonia-lyase |
| **OM** |  |  |  |  |
| PA1101 | -5.40447 | 2.27E-09 | *fliF* | flagellar MS-ring protein |
| PA1712 | -3.19963 | 3.2E-05 | *exsB* | exoenzyme S synthesis protein B |
| PA2800 | -2.70701 | 5.96E-05 | - | hypothetical protein |
| PA5107 | -2.21454 | 0.001957 | *Blc* | outer membrane lipoprotein Blc |
| **Cytoplasmic membrane** |  |  |  |  |
| PA4398 | -4.51676 | 1.35E-07 | - | two-component sensor |
| PA0375 | -4.2858 | 1.25E-08 | *ftsX* | cell division protein FtsX |
| PA4546 | -4.18337 | 1.14E-11 | *pilS* | two-component sensor PilS |
| PA1452 | -3.35969 | 1.25E-05 | *flhA* | flagellar biosynthesis protein FlhA |
| PA2235 | -3.29461 | 3.35E-05 | *pslE* | protein PslE |
| PA4910 | -3.10339 | 1.55E-06 | - | branched-chain amino acid ABC transporter ATP-binding protein |
| PA0016 | -3.03062 | 2.57E-07 | *trkA* | potassium transporter peripheral membrane protein |
| PA2873 | -3.00228 | 4.68E-08 | - | hypothetical protein |
| PA1703 | -2.99171 | 1.06E-05 | *pcrD* | type III secretory apparatus protein PcrD |
| PA0751 | -2.95272 | 5.24E-06 | - | hypothetical protein |
| **PA3210** | **-2.95042** | **3.15E-06** | *trkH* | **potassium uptake protein TrkH** |
| **PA5205** | **-2.87106** | **2.05E-11** | - | **multiple drug resistance protein MarC** |
| PA3489 | -2.78195 | 0.000644 | - | Na(+)-translocating NADH-quinone reductase subunit E |
| PA2604 | -2.73994 | 1.02E-08 | - | hypothetical protein |
| PA0890 | -2.66271 | 1.04E-07 | *aotM* | arginine/ornithine transporter AotM |
| PA5504 | -2.61934 | 0.000857 | - | D-methionine ABC transporter |
| PA3191 | -2.60578 | 0.000136 | - | two-component sensor |
| PA3211 | -2.60557 | 0.001151 | - | ABC transporter permease |
| PA1449 | -2.589 | 0.000294 | *flhB* | flagellar biosynthesis protein FlhB |
| PA3884 | -2.53839 | 3.88E-05 | - | hypothetical protein |
| PA1158 | -2.47781 | 0.001312 | - | two-component sensor |
| PA3102 | -2.46575 | 0.004995 | *xcpS* | general secretion pathway protein F |
| PA0392 | -2.40655 | 0.0011 | - | hypothetical protein |
| PA5256 | -2.38917 | 8.64E-07 | *dsbH* | disulfide bond formation protein |
| PA5043 | -2.38491 | 4.92E-05 | *pilN* | type 4 fimbrial biogenesis protein PilN |
| PA1691 | -2.34904 | 0.000369 | *pscT* | translocation protein in type III secretion |
| PA0184 | -2.34177 | 0.002138 | - | ABC transporter ATP-binding protein |
| PA4903 | -2.31684 | 8.63E-05 | - | major facilitator superfamily (MFS) transporter |
| PA5042 | -2.31445 | 0.00109 | *pilO* | type 4 fimbrial biogenesis protein PilO |
| PA2389 | -2.27523 | 4.17E-05 | *pvdR* | protein PvdR |
| PA4423 | -2.24433 | 0.000448 | - | hypothetical protein |
| PA2278 | -2.22646 | 0.001024 | *arsB* | arsenite-antimonite efflux pump ArsB |
| PA3963 | -2.21921 | 0.001037 | - | Transporter |
| **PA4954** | **-2.15623** | **0.001061** | ***motA*** | **flagellar motor protein MotA** |
| PA4332 | -2.13806 | 3.63E-06 | - | hypothetical protein |
| PA0753 | -2.12508 | 0.000215 | - | hypothetical protein |
| PA5258 | -2.11919 | 0.00121 | - | hypothetical protein |
| PA0374 | -2.10928 | 0.008412 | *ftsE* | cell division protein FtsE |
| PA3839 | -2.00907 | 0.007016 | - | sodium:sulfate symporter |
| PA2155 | -2.00863 | 0.007949 | - | cardiolipin synthase 2 |
| PA3857 | -2.00479 | 5.9E-05 | *pcs* | phosphatidylcholine synthase |
| **Extracellular** |  |  |  |  |
| PA1092 | -3.22535 | 9.38E-07 | *fliC* | flagellin type B |
| **PA1706** | **-2.54211** | **1.6E-05** | ***pcrV*** | **type III secretion protein PcrV** |
| PA4699 | -2.52982 | 0.00094 | - | hypothetical protein |
| PA4525 | -2.23285 | 0.000179 | *pilA* | type 4 fimbrial PilA |
| **PA0888** | **-2.12216** | **0.001585** | ***aotJ*** | **arginine/ornithine binding protein AotJ** |
| **Periplasm** |  |  |  |  |
| PA0766 | -5.29562 | 1.88E-09 | *mucD* | serine protease MucD |
| **PA1081** | **-3.3145** | **1.96E-05** | ***flgF*** | **flagellar basal body rod protein FlgF** |
| PA1092 | -3.22535 | 9.38E-07 | *fliC* | flagellin type B |
| PA4423 | -2.24433 | 0.000448 | - | hypothetical protein |
| **PA0888** | **-2.12216** | **0.001585** | ***aotJ*** | **arginine/ornithine binding protein AotJ** |
| **Cytoplasm** |  |  |  |  |
| PA5471 | -24.4276 | 3.21E-25 | - | hypothetical protein |
| PA0395 | -11.8779 | 5.47E-16 | *pilT* | twitching motility protein PilT |
| PA4547 | -8.10481 | 4.26E-17 | *pilR* | two-component response regulator PilR |
| PA1758 | -5.13388 | 2.96E-18 | *pabB* | para-aminobenzoate synthase component I |
| **PA1374** | **-4.67596** | **2.15E-08** | - | **hypothetical protein** |
| **PA1097** | **-4.59319** | **5.4E-12** | ***fleQ*** | **transcriptional regulator FleQ** |
| PA3824 | -4.29828 | 1.7E-08 | *queA* | S-adenosylmethionine--tRNA ribosyltransferase-isomerase |
| PA0396 | -4.02965 | 2.18E-08 | *pilU* | twitching motility protein PilU |
| PA5261 | -3.92591 | 8.22E-09 | *algR* | alginate biosynthesis regulatory protein AlgR |
| PA3192 | -3.78213 | 2.56E-06 | *gltR* | two-component response regulator GltR |
| PA4941 | -3.50556 | 2.91E-07 | *hflC* | protease subunit HflC |
| PA2235 | -3.29461 | 3.35E-05 | *pslE* | protein PslE |
| PA2043 | -3.27156 | 8.88E-05 | - | hypothetical protein |
| PA1572 | -3.21822 | 1.43E-06 | - | hypothetical protein |
| PA2260 | -3.13241 | 5.52E-05 | - | hypothetical protein |
| PA1110 | -3.05002 | 5.45E-06 | - | hypothetical protein |
| PA4524 | -3.0098 | 3.82E-05 | *nadC* | nicotinate-nucleotide pyrophosphorylase |
| PA5488 | -2.97215 | 3.64E-08 | - | hypothetical protein |
| PA3295 | -2.89537 | 1.31E-05 | - | HIT family protein |
| PA1987 | -2.86813 | 1.69E-05 | *pqqC* | pyrroloquinoline quinone biosynthesis protein PqqC |
| PA4942 | -2.80403 | 3.75E-05 | *hflK* | protease subunit HflK |
| **PA1455** | **-2.76169** | **3.83E-07** | *fliA* | **flagellar biosynthesis sigma factor** |
| PA3702 | -2.75103 | 0.000245 | *wspR* | two-component response regulator |
| **PA1127** | **-2.68943** | **0.000323** | - | **oxidoreductase** |
| PA2050 | -2.68555 | 0.000271 | - | RNA polymerase sigma factor |
| PA0338 | -2.66712 | 0.000282 | - | hypothetical protein |
| **PA3350** | **-2.6418** | **4.9E-05** | - | **flagellar basal body P-ring biosynthesis protein FlgA** |
| PA3070 | -2.52559 | 0.0002 | - | hypothetical protein |
| PA5493 | -2.51171 | 0.000288 | *polA* | DNA polymerase I |
| PA4231 | -2.49648 | 0.001317 | *pchA* | salicylate biosynthesis isochorismate synthase |
| PA3341 | -2.48126 | 0.00066 | - | transcriptional regulator |
| PA2550 | -2.42534 | 0.003606 | - | acyl-CoA dehydrogenase |
| PA4798 | -2.40225 | 0.000452 | - | hypothetical protein |
| PA3094 | -2.39923 | 6.39E-06 | - | transcriptional regulator |
| PA3945 | -2.3736 | 0.001944 | - | hypothetical protein |
| PA4191 | -2.34741 | 0.007044 | - | iron/ascorbate oxidoreductase |
| PA3723 | -2.30336 | 0.000115 | - | FMN oxidoreductase |
| PA2261 | -2.29666 | 0.002515 | - | 2-ketogluconate kinase |
| PA5115 | -2.27699 | 0.004989 | - | hypothetical protein |
| PA2255 | -2.2466 | 0.000223 | *pvcB* | paerucumarin biosynthesis protein PvcB |
| PA3444 | -2.21674 | 0.003829 | - | alkanesulfonate monooxygenase |
| PA1604 | -2.21501 | 0.001079 | - | hypothetical protein |
| PA3539 | -2.18164 | 2.68E-06 | - | hypothetical protein |
| PA3133 | -2.17678 | 0.000596 | - | transcriptional regulator |
| PA2254 | -2.17364 | 0.002019 | *pvcA* | paerucumarin biosynthesis protein PvcA |
| PA4902 | -2.15901 | 0.001438 | - | transcriptional regulator |
| PA2085 | -2.12973 | 0.004435 | - | ring-hydroxylating dioxygenase small subunit |
| PA2312 | -2.1194 | 0.007103 | - | transcriptional regulator |
| PA1950 | -2.11364 | 0.010999 | *rbsK* | Ribokinase |
| PA0413 | -2.10034 | 1.71E-06 | *chpA* | chemotactic signal transduction system protein |
| PA2496 | -2.10004 | 0.000213 | - | hypothetical protein |
| PA0937 | -2.08702 | 0.000629 | - | hypothetical protein |
| PA4508 | -2.07877 | 0.002013 | - | transcriptional regulator |
| PA3276 | -2.07523 | 0.00202 | - | hypothetical protein |
| PA4645 | -2.06194 | 0.007127 | - | hypoxanthine-guanine phosphoribosyltransferase |
| PA0893 | -2.05346 | 0.005301 | *argR* | transcriptional regulator ArgR |
| PA4617 | -2.02286 | 0.003027 | - | hypothetical protein |
| PA1186 | -2.02149 | 0.001298 | - | hypothetical protein |
| PA2003 | -2.01423 | 0.008875 | *bdhA* | 3-hydroxybutyrate dehydrogenase |
| PA2120 | -2.0132 | 0.003474 | - | hypothetical protein |
| PA5429 | -2.00956 | 0.001459 | *aspA* | aspartate ammonia-lyase |
| **Unknown** |  |  |  |  |
| **PA0271** | **-8.13811** | **2.14E-06** | **-** | **hypothetical protein** |
| PA3036 | -3.92635 | 9.6E-08 | - | hypothetical protein |
| PA4879 | -3.67382 | 2.31E-15 | - | hypothetical protein |
| **PA2559** | **-3.55392** | **2.38E-09** | **-** | **hypothetical protein** |
| PA3632 | -3.35329 | 1.46E-05 | - | hypothetical protein |
| PA4877 | -3.24691 | 8.73E-05 | - | hypothetical protein |
| PA5122 | -3.17151 | 1.62E-05 | - | hypothetical protein |
| PA2772a | -3.09495 | 8.4E-05 | - | hypothetical protein |
| **PA2009** | **-2.97886** | **2.88E-08** | ***hmgA*** | **homogentisate 1,2-dioxygenase** |
| **PA2238** | **-2.88138** | **5.37E-07** | ***pslH*** | **protein PslH** |
| PA4949 | -2.82889 | 0.000244 | - | hypothetical protein |
| PA3663 | -2.71457 | 0.001488 | - | hypothetical protein |
| PA1842 | -2.62859 | 0.000246 | - | hypothetical protein |
| PA3762 | -2.61944 | 0.000543 | - | hypothetical protein |
| PA4323 | -2.53315 | 0.000169 | - | hypothetical protein |
| PA1740 | -2.52962 | 5.72E-05 | - | hypothetical protein |
| PA3084 | -2.25805 | 0.001504 | - | hypothetical protein |
| PA3064 | -2.25203 | 1.25E-07 | *pelA* | PelA protein |
| PA4882 | -2.24131 | 0.000509 | - | hypothetical protein |
| PA4958 | -2.22137 | 0.001146 | - | hypothetical protein |
| PA1031 | -2.18395 | 0.005142 | - | hypothetical protein |
| PA5182 | -2.12966 | 0.000259 | - | hypothetical protein |
| PA2454 | -2.03603 | 0.002409 | - | hypothetical protein |
| PA3493 | -2.02273 | 0.008421 | - | hypothetical protein |
| PA5306 | -2.0162 | 0.003349 | - | hypothetical protein |

**Bold are common negatively selected genes in both conditions of infection.**

The cardiomyocyte infection condition without amikacin retains the entire population including adhered, internalized, released after host lysis, or actively released from the host cells unlike with amikacin which retains only the adhered and internalized. Among 143 conditionally essential genes, nineteen mutants are common in both infection conditions. Genes associated with flagella such as *flgE, flgF, flhA, flhB, fliA, fliC, fliF, motA*, and PA3350 are conditionally essential for cardiomyocyte infection in both conditions. *aotJ, aer, wbpJ, ponA*, *fleQ*, PA5205, *hmgA, trkH* and *pslH* are the common conditionally essential genes for infection. PA0271, PA1127, PA1374, PA2559 are the other common conditionally essential genes encoding the hypothetical protein. The reason for a few common conditionally essential genes for infection is unclear besides the actuality that the released pathogen can replicate without any constrain in antibiotic-free extracellular conditions. On the contrary, the released pathogen will be killed by amikacin present in the extracellular medium and only the host- protected pathogen will survive and proliferate. Other than the common conditionally essential genes, the crucial categories of the gene involved in type IV pili biosynthesis and regulation such as *pilA, pilT, pilU, pilO, pilR, pilS and pilN* and T3SS such as *pscT, pcrD, pcrV*, and *exsB*. These data suggest that the flagella and T3SS are essential for adhesion, invasion and host lysis / active release. And type IV pili are essential for host cell lysis or during the active release from the host. A recent study demonstrated that type IV pili are required by the toxins like ExlA to promote the contact between the pathogen and the host for further cytotoxic activities such as pore formation in the host cell membrane (Basso *et al*., 2017). Interestingly, we observed the requirement of three proteases during cardiomyocyte infection without antibiotics. *mucD, hflK*, and *hflC* are the proteases conditionally essential during cardiomyocyte infection which localize in OMVs. MucD is a serine protease and an important virulence factor of *Pseudomonas* in plant, insect, nematode and mammalian infections. It is associated with the production of an extracellular toxin to kill *C. elegans* (Yorgey *et al.*, 2001). Okuda *et al.*, (2011) demonstrated that the MucD is required by the toxin ExoS for IL-8 degradation to escape from the phagocytic killing by *P. aeruginosa*. Mochizuki *et al.*, (2014) investigated the role of MucD in *P. aeruginosa* during corneal infection. MucD suppresses the IL-1β, keratinocyte-derived cytokine and macrophage inflammatory protein-2 (MIP-2) and hence inhibits the neutrophil recruitment in the cornea during early infection. This facilitates *P. aeruginosa* to establish the infection by immune response evasion (Mochizuki *et al.*, 2014). *hflKC* is a two-gene member operon that encodes the accessory factors for FtsH protease. FtsH belongs to ATP-dependent zinc metalloprotease which is the major determinant of aminoglycoside resistance. HflK and HflC act as the regulators of FtsH and aids in the intrinsic aminoglycoside resistance of *P. aeruginosa* (Hinz *et al.*, 2011). Besides Aer, PonA, WbpJ, FlgE, and AotJ the other OMV localizing conditionally essential gene products are FtsX, PslE, PilA, AspA, PA4639, PA2800, PA4423, and PA5258. ArgR, FleQ, PA3341, PA3094, PA3133, PA4902, PA2312, and PA4508 are the conditionally essential transcriptional regulators during cardiomyocyte infection without amikacin.

#### 3.2.3. Conditionally enriched operon

Interestingly, a thirteen gene member operon that involves oxidative phosphorylation was found to be enriched during d-H9C2 infection. In both conditions of cardiomyocyte infection, most of the genes in *nuo* operon showed enhanced fitness. Table 3 provides the list of genes enriched in this operon with its fold change. Genes belong to this operon encode the subunits of type I NADH dehydrogenase which involves electron transfer to ubiquinone across the membrane by translocating the protons. Complex I play an important role in energy production. All the genes in this operon are essential for the production and assembly of the functional complex (Erhardt *et al.*, 2012). An earlier study demonstrated the inability of *nuo* mutants to grow anaerobically in *Pseudomonas aeruginosa* (Filiatrault *et al.*, 2006).

**Table 3.** List of gene members and their fold change in an enriched operon

| Locus Tag | Fold change |  | Gene | Product |
| --- | --- | --- | --- | --- |
|  | With amikacin | Without amikacin |  |  |
| **PA2637** | NA | 31.02858694 | *nuoA* | NADH dehydrogenase subunit A |
| **PA2638** | 7.99793686 | 11.70917298 | *nuoB* | NADH dehydrogenase subunit B |
| **PA2639** | 37.19144061 | 13.30317973 | *nuoD* | bifunctional NADH:ubiquinone oxidoreductase subunit C/D |
| **PA2640** | 3.160018015 | 11.12431574 | *nuoE* | NADH dehydrogenase subunit E |
| **PA2641** | 1.90736 | 11.65111592 | *nuoF* | NADH dehydrogenase I subunit F |
| **PA2642** | 3.105325153 | 14.98323237 | *nuoG* | NADH dehydrogenase subunit G |
| **PA2643** | 8.758440298 | 17.20452316 | *nuoH* | NADH dehydrogenase subunit H |
| **PA2644** | 23.12150498 | 12.82784012 | *nuoI* | NADH dehydrogenase subunit I |
| **PA2645** | NA | 15.56377344 | *nuoJ* | NADH dehydrogenase subunit J |
| **PA2646** | NA | 19.71054376 | *nuoK* | NADH dehydrogenase subunit K |
| **PA2647** | 47.59193344 | 8.956379216 | *nuoL* | NADH dehydrogenase subunit L |
| **PA2648** | 3.266331948 | 7.326796663 | *nuoM* | NADH dehydrogenase subunit M |
| **PA2649** | 12.42174022 | 6.606816739 | *nuoN* | NADH dehydrogenase subunit N |

Velmurugan *et al.*, (2007) revealed that the complex I is required for the inhibition of apoptosis during *Mycobacterium* infection. Particularly *nuo*G is the virulence gene that acts as an antiapoptotic factor. Our data shows the increased survival of the *nuo* gene mutants during cardiomyocyte infection. Since the *nuo* operon plays a crucial role in energy production, increased survival of its mutant during infection is a captivating outcome that needs further analysis.

### 3.3. Live-dead assay and CFU enumeration

Mutants of PA1561 (*aer*), PA5205 and *nuo*D were procured to validate the sequencing data. Figure 4a shows the representative images of the live-dead assay. In comparison with wild-type, PA1561::*lacZ* and PA5205::*phoA* showed lesser cytotoxicity (Figure 4b). PA2639::*phoA* showed significantly higher cytotoxicity in contrast with the wild-type. We found a significant (p<0.01) reduction in the number of CFU in both PA1561::*lacZ* (1E^+04^) and PA5205*::phoA* (5.7E^+05^) (Figure 4c). PA2639::*phoA* showed a significant (p<0.02) increased CFU/ml (2.35E^+08^) in contrast with PAO1 (1E^+07^). PA1561 encodes aerotaxis receptor Aer and PA5205 belong to MarC family of multiple-drug resistant proteins. One thousand-fold decrease of *aer* mutant and 100-fold decrease of PA5205*::phoA* in the intracellular load substantiates their conditional essentiality during cardiomyocyte infection. Ten-fold increased colonization of PA2639::*phoA* further validates the enhanced fitness of this gene after the disruption. PA2639 encodes NuoD- subunit of NADH: ubiquinone oxidoreductase. It appears that the cytotoxic effect of the mutants is related with their load during cardiomyocyte infection. Hence, the mutant showed enhanced fitness during infection resulted in severe cardiomyocyte death and vice versa.

**Figure 4.**
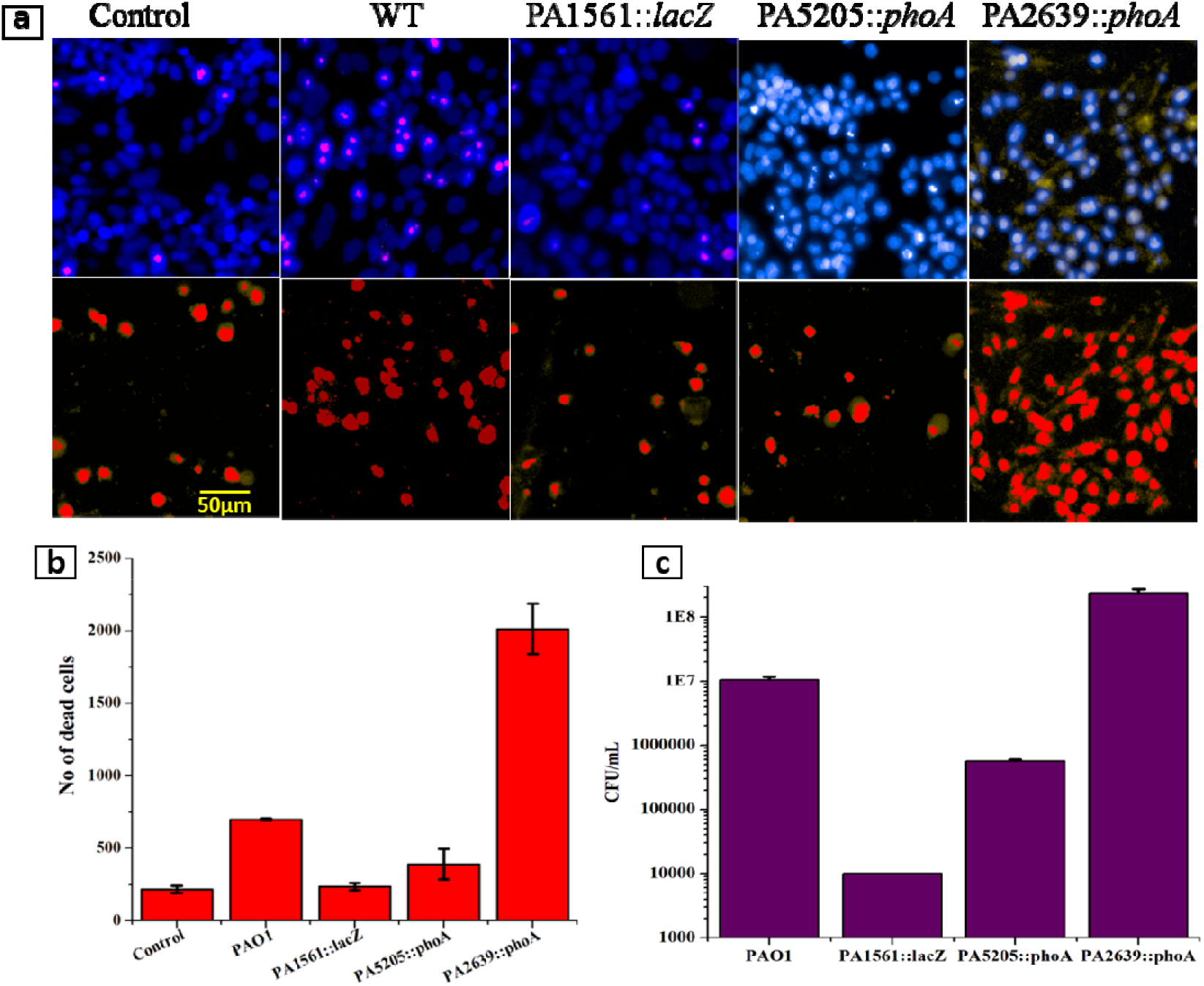
**a.** Representative images of the infected d-H9C2 cells with wild-type and mutants. **b.** The number of dead cells was quantified at 12 h of post-infection using PI stain. PI-stained cells were quantified using Harmony 3.0 software. c. CFU enumerations: d-H9C2 cells were infected with the mutants, and CFU was enumerated after 12 h of infection. The mean values of three independent experiments were shown in the bar chart.

## 4. Conclusion

The high incidence of *P. aeruginosa* especially in hospitals is due to the poor health status of the patients, the high prevalence of multidrug-resistant strains in hospitals, and the needless usage of broad-spectrum antibiotics. Biofilm formation, surface lipopolysaccharide, outer membrane proteins and secretion systems further enhance the pathogenesis. However, many efforts were taken in understanding the pathogenesis of *Pseudomonas* infection, better knowledge is obligatory for potential therapeutics to overcome the clinical burden due to *Pseudomonas*. Our findings aid in signifying the genetic requirements of *P.aeruginosa* during infection particularly at the intracellular-level. Further investigation of the core-set essential genes and conditional essential genes provide promising drug targets for this ubiquitous organism. Specifically, genes associated with flagella and T3SS are important for adhesion and invasion, while pili and proteases play a crucial role during host cell lysis. Furthermore, genes associated with amino acid transport & metabolism and signal transduction play a crucial role during stringent intracellular lifestyle whereas genes related to motility are important in the extracellular mode. OMVs play a significant role in the progression of the infection. Interestingly, the mutants of *nuo* operon are enriched in both conditions of infection and further study is needed to understand the molecular insights.

## Supporting information

Datafile-1

Datafile-2

Datafile-3

## Acknowledgments

Jothi Ranjani gratefully acknowledge DBT-MKU-IPLS and UGC-BSR for the financial support. She also acknowledges U.S. Vishnu for his technical support. The authors acknowledge UGC-NRCBS, DBT-IPLS, DST-PURSE and DST-FIST Programs of School of Biological Sciences, Madurai Kamaraj University.

