## Supplementary material for "Genome-wide identification of genetic requirements of *Pseudomonas aeruginosa* PAO1 for rat cardiomyocyte (H9C2) infection by insertion sequencing": Datafile-1

**Appendix-Table 1** List of strains and primers in this study

|  | **Strain** | **Characteristics** | **References** |
| --- | --- | --- | --- |
|  | ***P. aeruginosa* PAO1** | | |
|  | PW3811 | *aer::lacZ* | Jacobs *et al*., 2003 & Held et al., 2012 |
|  | PW9764 | *PA5205::phoA* | --do-- |
|  | PW5414 | *nuoD::phoA* | --do-- |
|  | ***E. coli*** | | |
|  | DH5α | F− Ø80lacZ M15 endA recA hsdR[rk−mk−] supE thi gyrA relAΔ[lacZYA-argF] U169 | Laboratory Stock |
|  | S17-1λ-pir | recA pro hsdR RP4-2-Tc::Mu-Km::Tn7λ-pir | de-Lorenzo *et al*, 1990 |
|  | **Plasmid**  pSAM_BT20 | ApR, GmR, transposon integration vector | Sivakumar *et al*., 2019 |
| **Primers** | | | |
| GENF | | 5’-TACTCGAGCGCGTCAATTCTCGAATTGACA-3’ | |
| GENR | | 5’- CAGCGGCCGCAGGGTTTTCCCAGTCAC-3’ | |
| TRANSF | | 5’-CACCGGATCCCAGTGTGATGGATTGACACATAG-3’ | |
| TRANSR | | 5’-TAGCGGCCGCAGGGTTTTCCCAGTCACG-3’ | |
| BioSamA* | | 5’-Biotin-TEG-CAAGCAGAAGACGGCATACGAAGACC-3’ | |
| LIB_PCR_5* | | 5’-CAAGCAGAAGACGGCATACGAAGACCGGGGACTTATCATCCAACCTGT-3’ | |
| LIB_PCR_3* | | 5’-AATGATACGGCGACCACCGAACACTCTTTCCCTACACGACGCTCTTCCGATCT-3’ | |

* - Goodman *et al*., 2011
